# Estrogen receptor alpha is required in GABAergic, but not glutamatergic, neurons to masculinize the brain

**DOI:** 10.1101/114835

**Authors:** Melody V. Wu, Jessica Tollkuhn

**Affiliations:** Cold Spring Harbor Laboratory, 1 Bungtown Road, Cold Spring Harbor, NY, 11724, USA

## Abstract

Masculinization of the rodent brain is driven by estrogen signaling during a perinatal critical period. Genetic deletion of estrogen receptor alpha (*Esr1/*ERα) results in altered hypothalamic-pituitary-gonadal (HPG) axis signaling and a dramatic reduction of male sexual and territorial behaviors. However, the requirement of ERα function in masculinizing distinct classes of neurons, and if these populations mediate components of male-typical behavior, remains unexplored. We deleted ERα in excitatory or inhibitory neurons using either a *Vglut2* or *Vgat* driver and assessed male behaviors. We find that *Vglut2*-Cre;*Esr1*^lox/lox^ mutant males lack ERa in the ventrolateral region of the ventromedial hypothalamus (VMHvl) and posterior ventral portion of the medial amygdala (MePV). These mutants recapitulate the increased serum testosterone levels seen with constitutive ERα deletion, but have none of the behavioral deficits. In contrast, *Vgat*-Cre;*Esr1*^lox/lox^ males with substantial ERα deletion in inhibitory neurons, including those of the principal nucleus of the bed nucleus of the stria terminalis (BNSTpr), posterior dorsal MeA (MePD), and medial preoptic area (MPOA) have normal testosterone levels, but display alterations in mating and territorial behaviors. These mutants also show demasculinized expression of androgen receptor (AR) and estrogen receptor beta (Esr2). Our results demonstrate that ERα masculinizes GABAergic neurons that gate the display of male-typical behaviors.

## Introduction

In mammals, females and males show differences in diverse social behaviors, including mating, aggression, and parental care. These behaviors are mediated by sexually dimorphic neural circuitry that develops under the control of gonadal hormones. In rodents, masculinization of the brain is driven primarily by estrogen signaling during a perinatal critical period. Circulating testosterone is converted to estradiol in the male brain by aromatase and this locally synthesized estrogen organizes the neural circuitry that regulates male-specific behaviors in adults (Amateau et al., 2004; Arnold, 2009; MacLusky and Naftolin, 1981; McCarthy, 2008; McCarthy et al., 2009; Naftolin and Ryan, 1975). Many of the effects of perinatal estradiol on sexual differentiation of the brain are exerted through estrogen receptor alpha (ERα). In males, genetic deletion of this receptor abolishes fertility, alters hypothalamic-gonadal-pituitary (HPG) axis signaling and impairs sexual and territorial behaviors (Ogawa et al., 1997; Scordalakes and Rissman, 2004; Wersinger et al., 1997), as well as social motivation (Imwalle et al., 2002) and social preference (Wersinger and Rissman, 2000). These innate behaviors are primarily regulated by hypothalamic regions that receive pheromonal chemosensory information from the accessory olfactory pathway. ERα is expressed in many of these areas including the medial amygdala (MeA), the principal nucleus of the bed nucleus of the stria terminalis (BNSTpr, referred to hereafter as BNST), the medial preoptic area (MPOA), and the ventrolateral region of the ventromedial hypothalamus (VMHvl) (Lee et al., 2014; Lin et al., 2011; Shughrue et al., 1997; Simerly et al., 1990). ERα-expressing neurons of the VMHvl and posterior ventral MeA (MePV) are largely glutamatergic (Hong et al., 2014;Lin et al., 2011; Sakurai et al., 2016; Ziegler et al., 2002), while ERα+ cells in the MPOA, BNST, and posterior dorsal MeA (MePD) are predominantly GABAergic (Cheong et al., 2015; Choi et al., 2005; Herbison and Fenelon, 1995; Hong et al., 2014; Hou et al., 2016; McHenry et al., 2017; Swanson and Petrovich, 1998; Unger et al., 2015). It has been proposed that the strong reciprocal GABAergic projections of the MeA and BNST generate “double-negative” input to their downstream hypothalamic regions, thereby gating the excitatory populations that drive behavioral displays (Choi et al., 2005; Hong et al., 2014; Swanson, 2000). However, the role of ERα itself in orchestrating sex differences in this hierarchy remains unexplored.

Previous studies have used viral delivery of shRNA against *Esr1* to assess the behavioral requirement for ERα in individual sexually dimorphic brain regions in adult and pubertal animals (Sano et al., 2016, 2013). However, there has been no analysis of male behaviors in mice with neural-specific deletions of the *Esr1* gene. The generation of cell-type-specific knock-in Cre drivers has made it possible to delineate the contribution of genetically-defined classes of neurons to specific parameters of mouse behaviors (Huang, 2014). We hypothesized that by deleting ERα in either glutamatergic or GABAergic neurons, we could begin to dissect the role of this receptor in regulating discrete components of male behavioral circuitry and physiology.

We crossed mice bearing a loxP-flanked allele of *Esr1* (*Esr1*^lox/lox^) (Correa et al., 2015; Feng et al., 2007) to either the *Slc17a6 (Vglut2)*-Cre or the *Slc32a1(Vgat)*-Cre knock-in mouse lines (Vong et al., 2011). To confirm loss of ERα expression during the perinatal critical period, we quantified ERα-expressing cells at p0 in the BNST, MePD, MePV, MPOA, and VMHvl of mutant and *Esr1*^lox/lox^ littermate control males. We measured serum testosterone, body weight, and seminal vesicles and testes weights in mutant males from both crosses as well as their littermates comprising the three control genotypes (wild-type, Cre only, and lox/lox only). Males of all eight genotypes were tested in assays for sexual behavior, inter-male aggression, and territory marking. Finally, *in situ* hybridization was used to investigate alterations in androgen receptor (*AR*) and estrogen receptor beta (*Esr2*) expression in adult brains from mutant males and *Esr1*^lox/lox^ littermate controls.

## Material and Methods

### Animals

*Vglut2*-Cre (Slc17a6tm2(cre)Lowl/J)(Vong et al., 2011) and *Vgat*-Cre (Slc32a1tm2(cre) Lowl/J) (Vong et al., 2011) mice were purchased from Jackson Labs and separately bred to *Esr1*^lox/lox^ mice(Feng et al., 2007). Mutant (*Vglut2*^Cre/+^;*Esr1*^lox/lox^ or *Vgat*^Cre/+^;*Esr1*^lox/lox^) and littermate control (*Vglut2/Vgat*^+/+^; *Esr1^+/+^* (wildtype), *Vglut2/Vgat*^Cre/+^; *Esr1*^+/+^ (Cre only), and *Vglut2/Vgat*^+/+^; *Esr1*^lox/lox^ (lox/lox only)) male mice were between 9 and 19 weeks of age at the start of behavioral assays. Histological assays were performed on mutant and respective lox/lox only littermate control male mice sacrificed on the day of birth (p0) or at >8 weeks of age (adult). 129SVE males from Taconic Farms and ovariectomized C57B1/6J females from Jackson Labs were purchased at 8 weeks of age to serve as stimulus animals for aggression and mating assays, respectively. Females were hormonally primed to be in estrus by injecting 10μg estradiol benzoate (EB) (Sigma E8515) 2 days, 5μg EB 1 day, and 500μg progesterone (Sigma P0130) 4-6 hours prior to mating assays. All stimulus animals were used in ≤ 8 assays and given ≥ 1 week recovery between assays.

All animals were maintained on a 12:12 light cycle (lights on at 01:00h) and provided food and water *ad libitum*. All procedures complied with NIH AALAC and CSHL IACUC guidelines.

### Behavioral assays

Animals from both the *Vgat*-Cre and *Vglut2*-Cre crosses were run side by side. All assays were performed ≥ 1 hour after the onset of the dark cycle. Animals were singly housed for 5 days, then tested twice for male mating, once for urine marking, and twice for residential aggression behaviors as previously described (Juntti et al., 2010; Wu et al., 2009). Fiji particle analysis software (NIH) was used to determine the number and size of urine spots for marking assays. Mating and residential aggression assays were videotaped under infrared illumination and scored offline using Observer software (Noldus). The experimenter was blinded to the genotype of the animals from the onset of single-housing to the completion of behavioral scoring. Mating assays were scored for mounting, intromission, ejaculation, and attacking (including bites, chasing, tumbling, and wrestling). Residential aggression assays were scored for attacking. The percentage of animals exhibiting each behavior in at least one assay was calculated. Latency to first display, total duration, and total frequency of each behavior, when performed, were averaged across the two assays.

Animals were sacrificed 3-5 days following the last behavioral assay. At the time of sacrifice, body weight was measured, blood was collected via submandibular bleed using a Goldenrod Lancet (MEDIpoint) into a Microtainer tube (BD 365963), then animals were anesthetized and perfused with phosphate buffered saline (PBS) followed by 4% paraformaldehyde. Testes, seminal vesicles, and brains were extracted following perfusion.

### Hormone assays

Collected blood was allowed to coagulate at room temperature for ≥30 minutes before centrifugation at 3000g for 10 minutes at 4°C. Resultant supernatant was transferred to a fresh Eppendorf tube and stored at -80°C. Serum testosterone was assayed in duplicate using a commercial ELISA kit (DRG EIA-1559) as directed. Standard curves were fit using 4PL online software available at mycurvefit.com. The DRG kit utilizes a monoclonal antibody and has a dynamic range between 0.083 and 16ng/mL. As provided by kit documentation, intra assay variance across an n of 20 is 4.16%, 3.28%, and 3.34% at low, mid, and high concentrations, respectively, while inter assay variance is 9.94%, 6.71%, and 4.73%.

### Histology

Brains were dissected from paraformaldehyde-perfused adult and freshly decapitated p0 animals. Brains were postfixed overnight then cryoprotected in 30% sucrose before being embedded in Shandon M-1 embedding matrix (Fisher 1310) and stored at -80°C. p0 brains were coronally cryosectioned at 20μm and serially adjacent sections were collected on two sets of slides and stored at -80°C. Adult brains were coronally cryosectioned at 60μm, collected on one set of slides, and processed immediately.

Immunolabeling of p0 brains was performed as previously described (Juntti et al., 2010; Wu et al., 2009). Antibodies used were rabbit anti-ERα (1:10K; EMD Millipore 06-935) and Cy3 donkey anti-rabbit (1:800, Jackson 711-165-152). In situ hybridization (ISH) against AR and Esr2 in adult brains was performed as previously described(Kurrasch et al., 2007;Wu et al., 2009). The probe to detect AR corresponds to bases 685-1724 of the cDNA and the probe to Esr2 corresponds to the sequence used by the Allen Brain Atlas (Lein et al., 2007).

Immunolabeled sections were imaged on an LSM 710 confocal microscope at 20x magnification. The center optical slice was imaged from one ROI for each 20μm section spanning the entire antero-posterior extent of both sides of the BNST, MPOA, and MePD of *Vgat*-Cre;*Esr1*^lox/lox^ mutant and control animals, and of the VMHvl and MePV of *Vglut2*-Cre;*Esr1*^lox/lox^ mutant and control animals. Identification of the p0 MePV was aided by visual comparison to prenatal Lhx9 expression (Garcia-Lopez et al., 2008). Multiple slices were tiled together to obtain representative images for Figure 1. The number of ERα+ cells was quantified using Fiji analysis software (NIH) with experimenters blind to animal genotype. Briefly, the ROI was outlined in each image, the image was made binary, Gaussian blur with sigma=2 was applied, the image was made binary again, watershed segmentation was applied, and particle analysis was performed with size set to 20-infinity and circularity set to 0-1. As immunolabeling was performed on every other section, the number of counted particles was multiplied by 2 to obtain total numbers of ERα+ cells.

**Figure 1:**
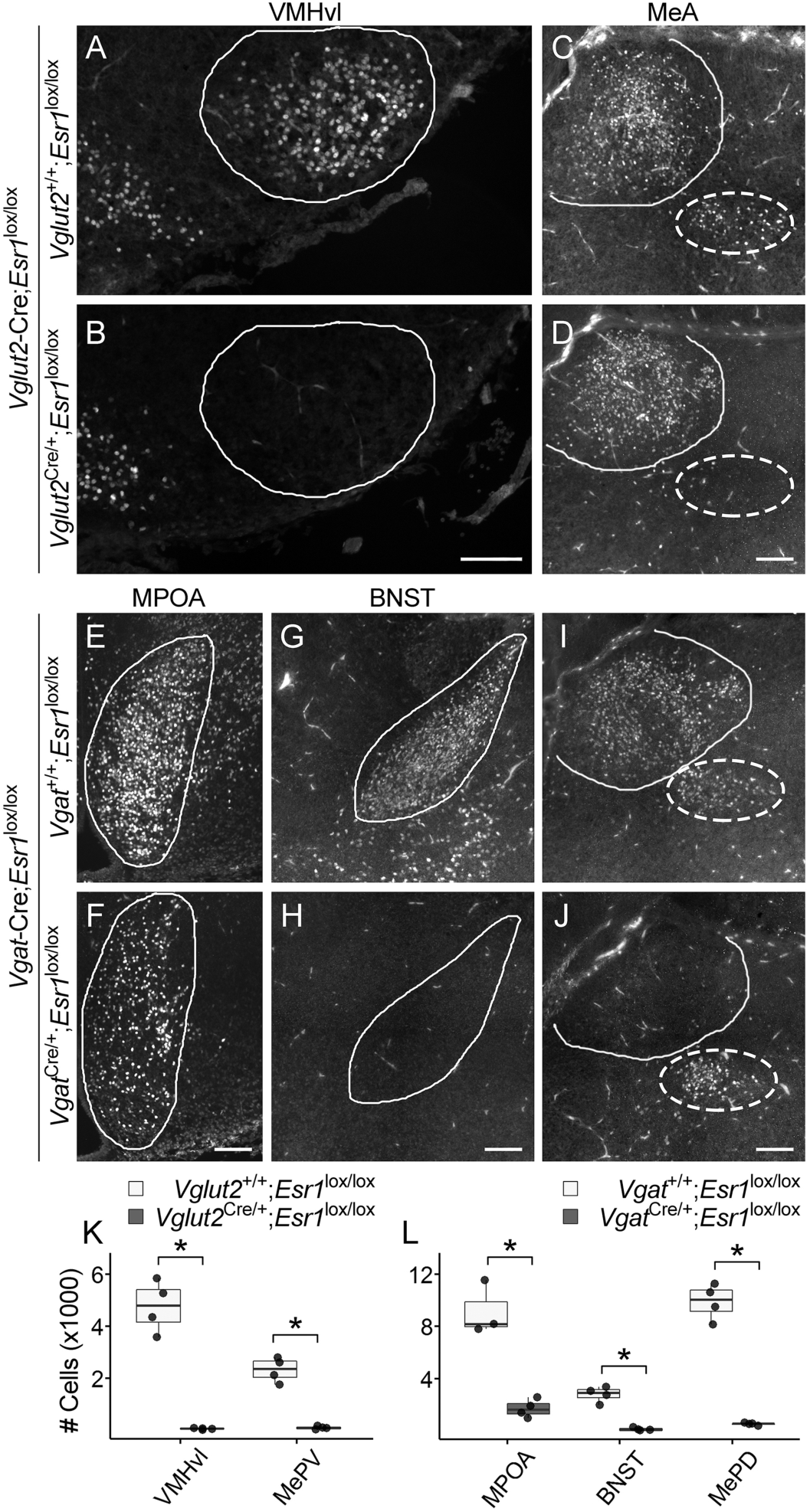
Neonatal deletion of *Esr*1 in *Vglut2*+ and *Vgat*+ neurons. Shown are representative images of ERα expression in the VMHvl (**A,B**) and MeA (**C,D**) of lox/lox only control and *Vglut2*-Cre;*Esr1*^lox/lox^ mutant male littermates taken on the day of birth (p0). Representative images of the MPOA (**E,F**), BNST (**G,H**), and MeA (**I,J**) in lox/lox control and *Vgat*-Cre;*Esr1*^lox/lox^ mutant p0 male littermates are also depicted. Solid lines outline the VMHvl (**A-B**), MPOA (**E-F**), BNST (**G-H**) and MePD (**C,D,I,J**) and dotted lines outline the MePV (**C,D,I,J**). ERα expression is virtually absent in the VMHvl and MePV of *Vglut2*-Cre mutants (**K**) and in the BNST and MePD of *Vgat*-Cre mutants (**L**). Scalebars = 100μ. Boxplots denote median and 1^st^ and 3^rd^ quartiles. Whiskers denote 1.5^*^ interquartile range. *p<0.05, Mann Whitney U test.

ISH processed sections were imaged on a Zeiss Observer inverted microscope at 5x magnification under brightfield illumination. Each 60μm section was imaged, spanning the entire antero-posterior extent of the BNST and MePD. A standardized mask for each section of each region was used to outline the ROI as well as a background region for each image. Utilizing Fiji analysis software (NIH), each image was color inverted, the mean intensity value of the background region was subtracted from the ROI, expression levels were normalized, and the mean intensity of the resultant ROI was calculated. Average intensity for each section was then summed for a total region intensity.

### Statistics

Fisher’s exact test with a 2×4 contingency table was used to analyze categorical data across all groups. For α=0.05, subsequent post-hoc tests using 2×2 tables and Bonferroni correction were carried out. Differences were deemed significant for p<0.017 (0.05/3). Non-parametric tests were used for all other analyses due to non-normality of datasets(Krzywinski and Altman, 2014). Behavioral and physiological data was analyzed using the Kruskal-Wallis omnibus test followed by post-hoc analysis as necessary (α=0.05) with Dunn’s test for multiple comparisons with one control as provided by the PMCMR R package (Pohlert, 2016). For post-hoc measurements, wildtype, lox/lox only, and Cre only control littermates were each compared against mutant animals. Significance was assigned at p<0.05. Histological data was analyzed using the Mann-Whitney U test. The full panel of statistical results is provided in Table S1.

## Results

### Deletion of *Esr1* in excitatory or inhibitory neurons is complete at p0

We assessed ERα expression in order to determine the extent of *Esr1* deletion at p0, the time of the perinatal testosterone surge in mice (Motelica-Heino et al., 1993). We dissected brains from mutant males (*Vglut2*-Cre;*Esr1*^lox/lox^ or *Vgat*-Cre;*Esr1*^lox/lox^, n=4) and their respective control littermates lacking Cre (*Vglut2^+/+^;Esr1*^lox/lox^ or *Vgat*^+/+^;*Esr1*^lox/lox^, n=4) on p0, and quantified ERα+ cells in MPOA, BNST, MePD, MePV, and VMHvl. We find that ERα is almost entirely absent from the VMHvl and MePV of *Vglut2*-Cre;*Esr1*^lox/lox^ males (Fig 1A-D,K), but not changed in the MPOA, BNST, and MePD (Fig S1E-H and Fig1C-D). In *Vgat*-Cre;*Esr1*^lox/lox^ males, ERα is deleted in >96% of the BNST, 94% of the MePD and 82% of the MPOA (Fig 1E-J,L) (U=16, p=0.021 for all comparisons) and is unaffected in the VMHvl and MePV (Fig S1A-B and Fig1I-J). We also observed a reduction in cortical ERα in *Vglut2,* but not *Vgat* mutant animals, however we did not perform cell counts throughout the cortex (Fig S1 C,D,I,J). Our results confirm that deletion of *Esr1* in targeted cells is already complete at birth.

### Deletion of *Esr1* in excitatory or inhibitory neurons does not result in gross physiological deficits

We hypothesized that males with neural-specific deletions of Esr1 would have largely normal physiology as ERα expression is retained in peripheral tissues, such as the gonads, the pituitary and white adipose tissue. To assess the general physiology of animals lacking *Esr1* in *Vglut2*+ excitatory neurons or all inhibitory neurons, we examined body weight, testes weight, serum testosterone levels, and seminal vesicle weight in *Vglut2*-Cre;*Esr1*^lox/lox^ mutants (n=20) and their littermate controls—wildtype (n=5), lox/lox only (n=10), and *Vglut2*-Cre only (n=10). We measured the same parameters in *Vgat*-Cre;*Esr1*^lox/lox^ mutants (n=16) and their littermate controls—wildtype (n=7), lox/lox only (n=9), and *Vgat*-Cre only (n=8). *Vglut2*-Cre;*Esr1*^lox/lox^ mutant animals exhibited normal body and testes weights (Fig 2A-B). However, deletion of *Esr1* in *Vglut2*+ cells resulted in significantly increased serum testosterone levels (H=15.68, d.f.=3, p=0.0013) as compared with both wildtype and *Vglut2*-Cre only control animals (Dunn’s post-hoc p=0.045, p=0.0011 respectively) (Fig 2C). Seminal vesicle weight, a readout of circulating testosterone levels, was also higher in mutants compared to *Vglut2*-Cre only control animals (H=8.60, d.f.=3, p=0.035; Dunn’s post-hoc p=0.035) (Fig 2D) as observed in constitutive ERαKO males (Rissman et al., 1997). Male mice with a constitutive deletion of *Esr1* (ERαKOs) display increased body weight (Heine et al., 2000), decreased testes weight, elevated serum testosterone levels, and infertility (Eddy et al., 1996; Rissman et al., 1997). These results demonstrate that ERα is required in excitatory neurons to provide negative HPG feedback in males.

**Figure 2:**
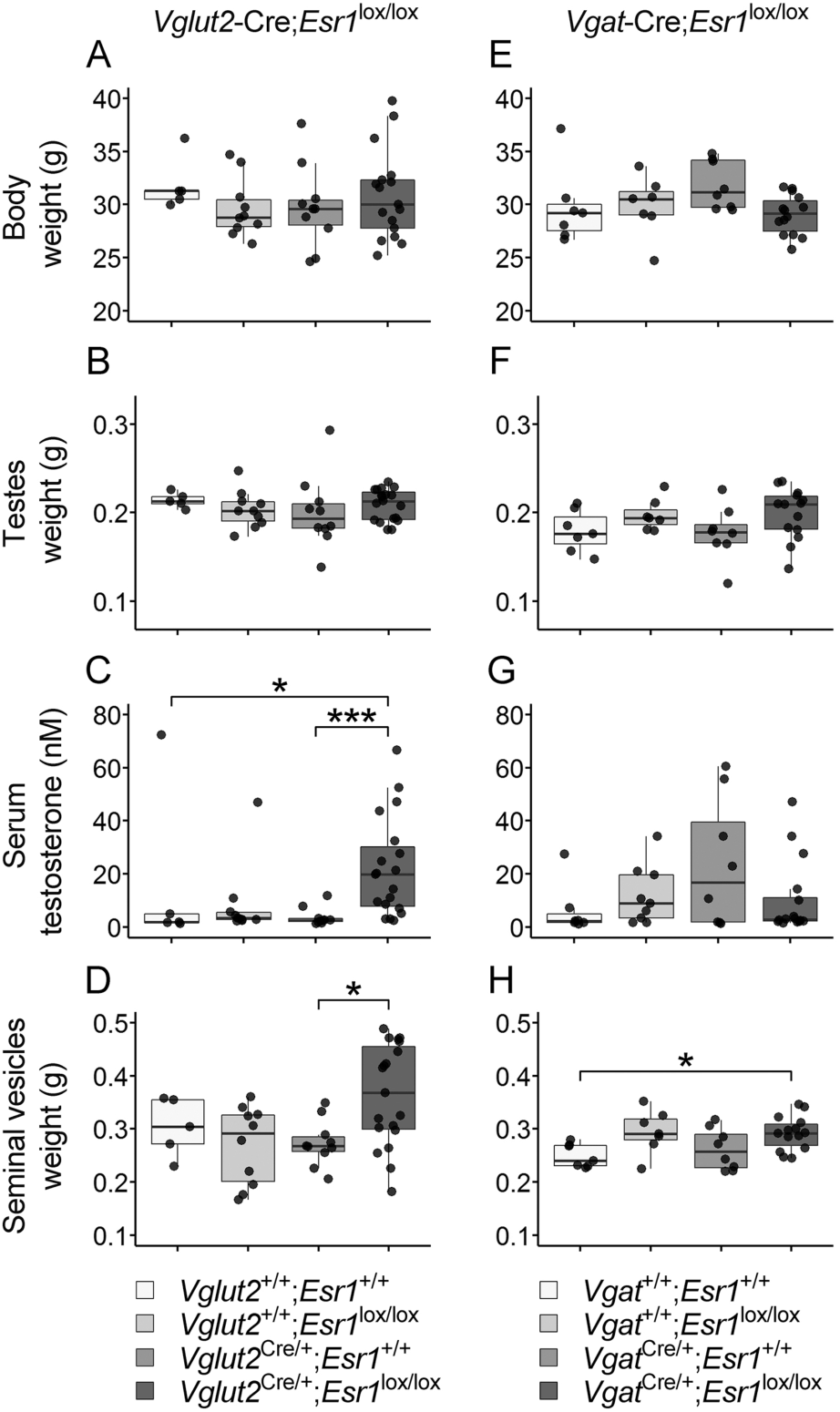
Physiological effects of *Esr1* deletion in *Vglut2+* and *Vgat+* neurons. Body weight (**A,E**), testes weight (**B,F**), serum testosterone levels (**C,G**), and seminal vesicles weight (**D,H**) are shown for *Vglut2*-Cre;*Esr1*^lox/lox^ mutant and littermate controls (**A-D**) and *Vgat-Cre;Esr1*^loxl/lox^ mutant and littermate controls (**E-H**). Boxplots denote median and 1^st^ and 3^rd^ quartiles. Whiskers denote 1.5^*^ interquartile range. *p<0.05, ***p<0.005, Kruskal Wallis omnibus test followed by Dunn’s post-hoc test for multiple comparisons with one control.

By contrast, deletion of *Esr1* in *Vgat*+ cells did not result in physiological deficits. Though seminal vesicles of mutant animals were slightly heavier than wildtype controls (H=9.48, d.f.=3, p=0.024; Dunn’s post-hoc p=0.030) (Fig 2H), body weight, testes weight, and serum testosterone levels were indistinguishable between mutant and control animals (Fig 2E-G). These changes do not appear to affect fertility as, in contrast to *Vgat*-Cre;*Esr1*^lox/lox^ and *Vglut2*-Cre;*Esr1*^lox/lox^ mutant females (Cheong et al., 2015), mutant males consistently produce litters (data not shown). Thus, the reproductive and neuroendocrine roles of ERα in males can be dissociated from other neural phenotypes by deletion of *Esr1* in genetically-defined populations of neurons.

### *Esr1* expression is not necessary in *Vglut2*+ neurons for male-typical behaviors

Vglut2-Cre is expressed in excitatory neurons throughout the brain, with enrichment predominantly in subcortical regions including the thalamus, hypothalamus and amygdala, as well as the piriform cortex (Cheong et al., 2015; Vong et al., 2011). We hypothesized that deletion of Esr1 in Vglut2+ neurons would result in decreased displays of male sexual and territorial behaviors. Notably these mutant males lack ERα in the VMHvl (Figure 1A-B,K), a region which drives the display of both mating and inter-male aggression when stimulated (Falkner et al., 2016; Flanagan-Cato et al., 2006; Lin et al., 2011; Yang et al., 2013). Surprisingly, *Vglut2*-Cre;*Esr1*^lox/lox^ mutant males display wildtype-typical male mating, urine marking, and resident aggression. The percent of animals displaying each component of mating and aggressive behavior is indistinguishable from control littermates (Fig 3A-E), and when displayed, there is no difference in the frequency or duration of these behaviors (Fig S2). In addition, both the number of urine spots deposited and the total area covered by the spots are comparable between mutant and control animals (Fig 3F-K). These results suggest that *Esr1* expression in VGlut2+ excitatory neurons is dispensable for male sexual and territorial behaviors.

**Figure 3:**
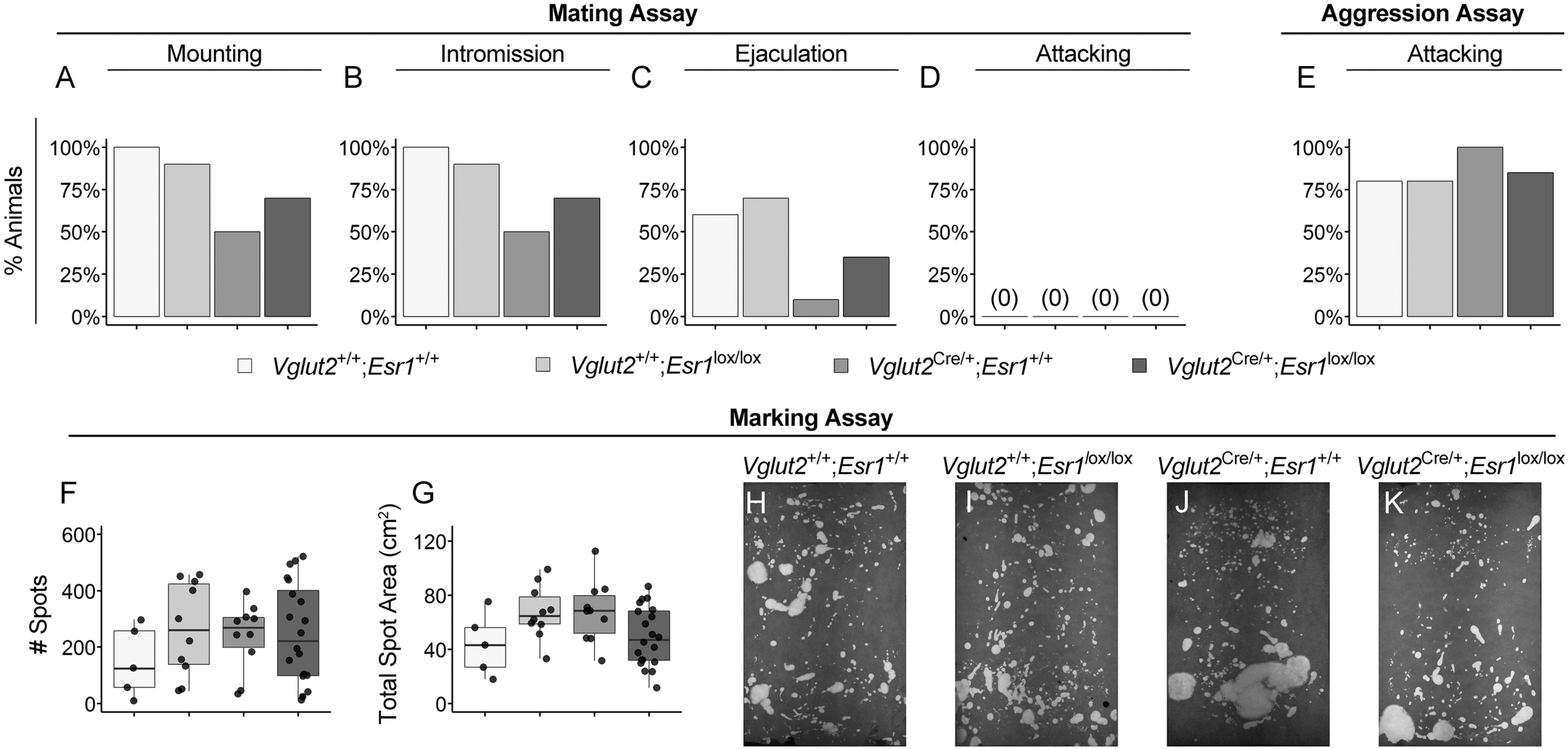
No behavioral effect of *Esr1* deletion in *Vglut2*+ neurons. Depicted are the percentages of animals that display mounting (**A**), intromission (**B**), ejaculation (**C**), and attacking behavior (**D**) in at least one out of two mating assays and the percentage of animals that display attacking behavior (**E**) in at least one out of two aggression assays. Number of spots deposited (**F**) and total urine coverage area in a marking assay (**G**) are shown alongside representative urine marking assays (**H-K**). Boxplots denote median and 1^st^ and 3^rd^ quartiles. Whiskers denote 1.5^*^ interquartile range.

### Deletion of *Esr1* in inhibitory neurons dysregulates male mating and demasculinizes territorial marking

We next examined sex-typical behaviors in *Vgat*-Cre;*Esr1*^lox/lox^ mutant males, which lack ERα in the BNST, MePD, and much of the MPOA (Fig 1E-J, L). The percentage of animals displaying mounting and intromission in at least one of two 30 minute mating assays did not differ between mutant mice and control littermates (Fig 4A-B). However, there was a significant decrease in the percentage of mutant animals that ejaculated within the assay(Fisher’s Exact 2×4 p=0.0002) compared to both lox/lox only and *Vgat*-Cre only control littermates (Fisher’s Exact 2×2 p=0.009, p=0.0002 respectively) (Fig 4C). The quality of most mating behaviors is not changed by *Esr1* deletion as the frequency, total duration, and latency to perform mounts or intromits, and even the latency to ejaculate, when displayed, were equivalent between mutants and control littermates (Fig S3).

**Figure 4:**
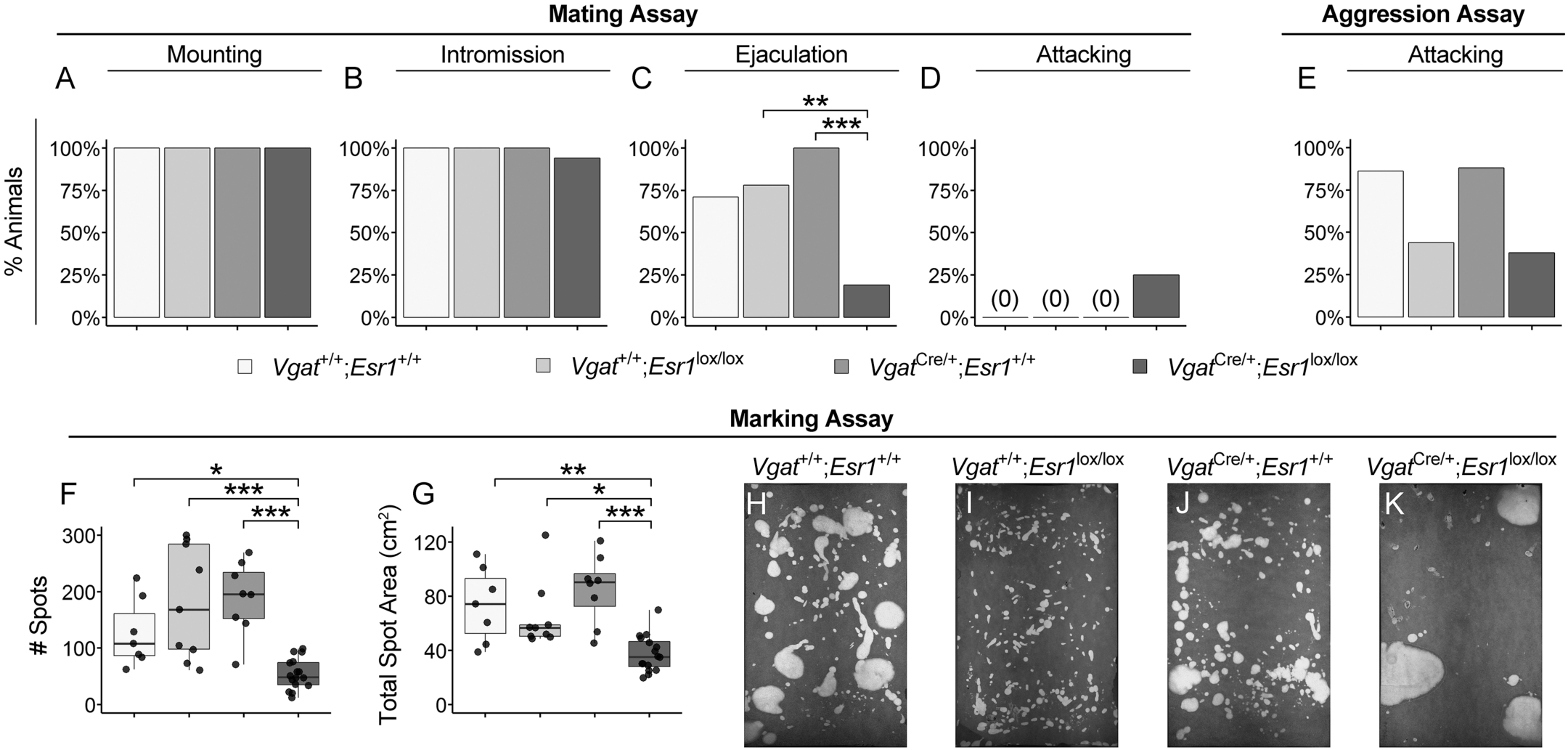
Deletion of *Esr1* in *Vgat+* cells results in deficits in mating and marking behavior. Depicted are the percentages of animals that display mounting (**A**), intromission (**B**), ejaculation (**C**), and attacking (**D**) behavior in at least one out of two mating assays and the percentage of animals that display attacking behavior (**E**) in at least one out of two aggression assays. Number of spots deposited (**F**) and total urine coverage area in a marking assay (**G**) are shown alongside representative urine marking assays (**H-K**). Boxplots denote median and 1^st^ and 3^rd^ quartiles. Whiskers denote 1.5^*^ interquartile range. **p<0.01, ***p<0.005, Fisher’s 2×4 contingency table followed by post-hoc Fisher’s 2×2 contingency table with Bonferroni correction (A-E). *p<0.05, **p<0.01, ***p<0.005, Kruskal Wallis omnibus test followed by Dunn’s post-hoc test for multiple comparisons with one control (F-G).

Strikingly, 25% of *Vgat*-Cre;*Esr1*^lox/lox^ mutant males attacked estrus females in mating assays, a phenomenon never seen in control animals (Fig 4D). All of the animals that attacked females did display male-typical mating behavior, with 2 out of 4 mutants displaying both intromissions and attacks in the same assay. However, these mutants do not simply exhibit increased, indiscriminate, attack behavior, as there is no increase in the percentage of animals attacking intruder males in a resident aggression assay compared to littermate controls (Fig 4E). The frequency and total duration of attacks, as well as the latency to attack intruder males, was not different in mutant animals (Fig S3).

When placed in a novel environment, isolated wildtype males typically deposit many small urine spots spanning the length and width of the environment as a manifestation of territorial marking (Desjardins et al., 1973). This behavior can be elicited in castrated adult males by either testosterone or estrogen (Kimura and Hagiwara, 1985; Nyby, 1992), while nervous system-specific AR mutant mice do not mark (Juntti et al., 2010). However, a masculinized urine marking pattern can be elicited in females treated with estradiol postnatally(Wu et al., 2009), suggesting that masculinization of marking behavior may also be downstream of ERα signaling. Strikingly, *Vgat*-Cre;*Esr1*^lox/lox^ mutant mice display profoundly demasculinized territorial marking behavior. Mutant males deposit three-fold fewer urine spots (H=21.97, d.f.=3, p=0.00007) compared to each of their littermate controls (Dunn’s post-hoc p=0.037, p=0.0006, p=0.0003 against wildtype, lox/lox only, and *Vgat*-Cre only control littermates respectively) (Fig 4F). This deficit is also reflected in a significant drop in total area covered by urine (H=20.45, d.f.=3, p=0.0001; Dunn’s post-hoc p=0.008, p=0.012, p=0.0002 against wildtype, lox/lox only, and *Vgat*-Cre only control littermates respectively) (Fig 4G). Finally, urine deposited by mutants was pooled in the corners of the cage (Figure 4K), rather than spread over the cage floor (Figure 4H-J). Taken together, these results demonstrate that ERα acts in GABAergic neurons to masculinize discrete components of male behavior.

### Deletion of *Esr1* in *Vgat+* cells demasculinizes gene expression

Masculinization of sexually dimorphic behavior is known to be critically dependent on the male perinatal surge of testosterone, and its subsequent aromatization to estradiol (Motelica-Heino et al., 1993; Wu et al., 2009). In addition to masculinization of behavior, such early estrogen signaling has long-term effects on gene expression (Nugent et al., 2015; Ratnu et al., 2017; Xu et al., 2002, 2012), including setting up dimorphic expression of AR in the BNST and MPOA (Juntti et al., 2010). Esr2 expression also appears to be under the control of early estrogen signaling as its mRNA expression is downregulated in postnatal female rats given estradiol at birth (Cao et al., 2012) while ERβ protein levels are upregulated in the BNST of adult systemic ERαKO male mice (Nomura et al., 2003). We thus hypothesized that our *Vgat*-Cre;*Esr1*^lox/lox^ mutant males might exhibit demasculinization of AR and Esr2 expression. Indeed, expression of AR mRNA is significantly lower in both the BNST and MePD of *Vgat*-Cre;*Esr1*^lox/lox^ adult mutants compared to littermate controls (U=15, p=0.043 for both comparisons) (Fig 5A-D,I). While there appears to be no change in Esr2 expression in the MePD (U=8, p>0.99), there is a significant increase in its expression in the BNST of mutant animals (U=15, p=0.043) (Fig 5E-H,J), <u>suggesting the possibility of a compensatory mechanism or mutually-balancing function via ERβ signaling in this region</u>. AR is expressed robustly in the VMHvl and the MePV (DonCarlos et al., 1995), yet we did not observe any change in AR expression in *Vglut2*-Cre;*Esr1*^lox/lox^ males (data not shown). Taken together, our results demonstrate that ERα regulates Esr2 and AR expression in specific populations of *Vgat+* neurons.

**Figure 5:**
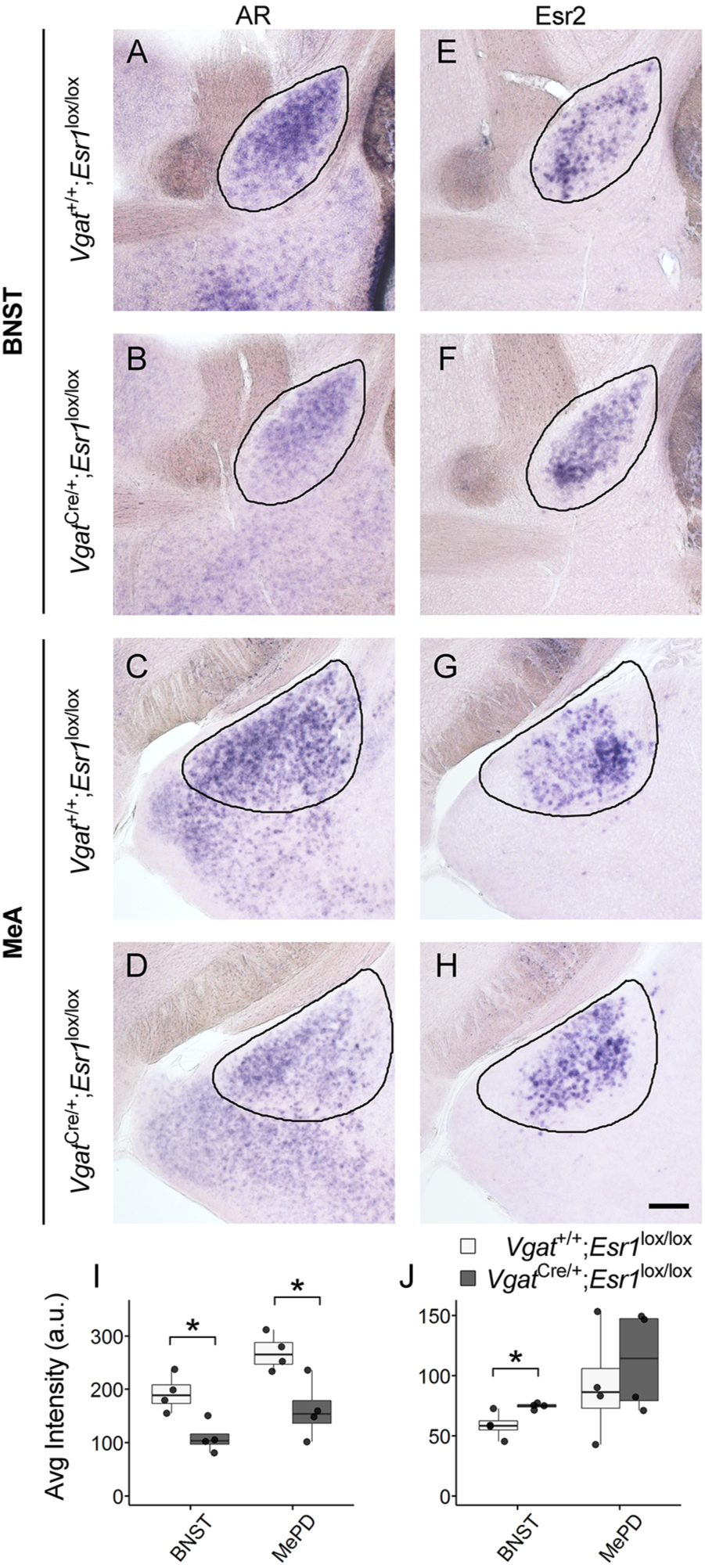
Demasculinized AR and Esr2 expression in *Vgat*-Cre;*Esr1*^lox/lox^ males. Representative images are shown of AR (**A-D**) and Esr2 (**E-H**) expression in the BNST (**A,B,E,F**) and MeA (**C,D,G,H**) of *Vgat*-Cre;*Esr1*^lox/lox^ mutant and littermate control adult males. Black lines outline ROIs of expression intensity quantification. Average intensity of *AR* mRNA expression is significantly downregulated in mutants (**I**) while *Esr2* mRNA expression is increased in the BNST of mutant males (**J**). Scalebar = 200μ. Boxplots denote median and 1^st^ and 3^rd^ quartiles. Whiskers denote 1.5^*^ interquartile range. *p<0.05, Mann Whitney U test.

## Discussion

### Male behaviors do not require ERα in *Vglut2*+ excitatory neurons

Females and males show distinct behavioral responses to novel males: only females can be sexually receptive and males generally display territorial aggression. These sexually dimorphic behaviors are regulated by shared neural circuitry between the two sexes, while both sexes require ERα for correct sexual differentiation of this circuitry. Our goal in this study was to begin to parse how ERα function specifies the flow of information from pheromonal input to male-typical behavioral output. Unexpectedly, we find that loss of ERα protein in glutamatergic neurons enriched in subcortical structures has no effect on male sexual behavior, aggression, or territory marking; mutant males show wild-type levels of every behavioral parameter tested. Mutant males do possess increased levels of serum testosterone and increased seminal vesicle weights, indicating that HPG feedback is disrupted in these animals, as demonstrated previously in *Vglut2*-Cre;*Esr1*^lox/lox^ mutant females and constitutive ERaKO males (Cheong et al., 2015; Rissman et al., 1997). ERα protein is almost entirely absent from the MePV and VMHvl of mutant males at birth (Figure 1K), demonstrating that ERα is dispensable for perinatal organization of these brain regions. Previous studies have demonstrated that ERα+ neurons in the VMHvl regulate aggression and mating behavior in males. Optogenetic activation of ERα+ neurons in male VMHvl induces time-locked attacks, and can also induce mounting behavior (Lee et al., 2014). Ablating progesterone-receptor (PR)-expressing cells that are also ERα+ in adult males results in reduced mounting and intromission toward females and decreased attacks towards males (Yang et al., 2013). Our results show that ERα itself is not required in VMHvl neurons for the display of male mating and aggression, and suggests that masculinzation of circuits downstream of ERα+-VMHvl neurons is necessary for VMH-elicted aggression. These results contrast with findings in females, where virally-mediated shRNA knockdown of ERα in VMHvl decreases both receptive behavior, sexual motivation, and PR expression (Musatov et al., 2006; Spiteri et al., 2010). Estrogens also increase neural activity of female VMHvl neurons, both in slice (Kow et al., 2005; Kow and Pfaff, 1985) and in estrous females interacting with males (Nomoto and Lima, 2015). Finally, many labs have demonstrated that ovarian hormones alter synaptic connectivity and spine number in females (reviewed in Flanagan-Cato, 2011) and that there are estradiol dependent sex differences in synaptic structure and organization in the VMH (Schwarz et al., 2008; Todd et al., 2007). Nevertheless, the complete absence of a behavioral phenotype in males lacking ERα in glutamatergic neurons suggests that the predominant organizational role of this receptor in male-typical behaviors is in inhibitory neurons that modulate the development or activity of the VMHvl.

### ERα expression in inhibitory neurons regulates male sexual and territorial behaviors

We find that loss of ERα in GABAergic neurons alters both mating and territorial behaviors (Fig 4), but does not affect circulating hormone levels or body weight (Fig 2). At birth, *Vgat*-Cre;*Esr1*^lox/lox^ males lack ERα expression in the BNST and MePD, and there is a profound decrease in ERα+ cells in the MPOA (Fig 1L). The MPOA is a crucial regulator of male sexual behavior (Arendash and Gorski, 1983; Christensen et al., 1977; Edwards and Einhorn, 1986; Hull and Dominguez, 2007) and accordingly, only 25% of *Vgat*-Cre;*Esr1*^lox/lox^ mutant males ejaculated in a 30 minute assay, compared to 70-95% ejaculation in control genotypes (Fig 4). All mutants display normal numbers of mounts and intromits, consitent with initial studies for gonadally intact ERαKO males (Ogawa et al., 1997). Indeed treatment of constitutive ERαKO mutants with a dopamine agonist restores wild-type levels of mounting and intromission, demonstrating that ERα is not required for the development of circuitry that underlies male sexual behavior (Wersinger and Rissman, 2000). 25% of mutant males attacked females in a mating assay, at times attacking and mounting in the same assay. This aberrant display of aggression towards females was first observed in ERαKO males. These animals also have deficits in social recognition (Imwalle et al., 2002) and do not show a social preference for investigating females compared to males (Scordalakes and Rissman, 2003). The medial amygdala displays sexually dimorphic responses to olfactory cues from males or females; neurons in the male MeA are more selective for female odors, compared to male or predator odors (Bergan et al., 2014). This male bias for female odors is blunted in males mutant for aromatase, which is primarily expressed in the BNST and MePD (Wu et al., 2009). Our results suggest that male-typical recognition and processing of pheromonal cues requires ERα in the BNST and MePD.

*Vgat*-Cre;*Esr1*^lox/lox^ males show deficits in male-typical territorial behaviors, with reduced inter-male aggression and demasculinized territory marking. Mutants do not urine mark the cage floor, but pool their urine in the corners of the cage in the pattern of wild-type females or subordinate males. Previous work implicated estrogen as the master regulator of male territorial behavior circuitry. Treating wild-type females with estradiol in the first two weeks of life leads to a masculinized pattern of urine-marking (Wu et al., 2009) and males with a neural deletion of AR mark in a wild-type pattern, but with reduced intensity (Juntti et al., 2010). Here, we demonstrate that male territory-marking behavior requires ERα expression specifically in inhibitory neurons. We attribute the marking phenotype specifically to decreased expression of ERα in GABAergic neurons of the MPOA. Chemogenetic inhibition of these neurons in dominant males produces a subordinate urine-marking pattern through disinhibition of Crh+ neurons in the pontine micturition center (PMC)(Hou et al., 2016). Our experiments suggest that ERα signaling masculinizes MPOA control of urine marking pattern and intensity. *Vgat*+ MPOA neurons also promote social reward in both sexes and this is enhanced by estrogen in females(McHenry et al., 2017). It would be interesting to assess additional social behaviors, such as parental behavior, in *Vgat*-Cre;*Esr1*^lox/lox^ mutants of both sexes.

### Gene expression changes in demasculinized *Vgat*-Cre;*Esr1*^lox/lox^ mutant males

ERα is a nuclear receptor transcription factor that activates gene expression in the presence of its ligand estrogen. We find that loss of ERα in GABAergic neurons leads to decreased AR expression in the BNST and MePD. Our results are consistent with earlier studies demonstrating that perinatal estrogen increases the number of AR-expressing cells in males (Juntti et al., 2010), and that ERαKO mutants have decreased AR expression (Wersinger et al., 1997). Perinatal estrogen is also known to masculinize the number of neurons in the BNST (Hisasue et al., 2010; Wu et al., 2009) and, as we did not compare the number of neurons between wildtype males, mutant males, and wildtype females, it is therefore possible that the decreased AR expression is due to decreased cell survival in mutant males rather than direct changes in the levels of AR gene activation in individual cells. However, we also find that Esr2 is increased in the BNST of mutant males in adulthood, as shown previously in constitutive ERαKOs (Nomura et al., 2003) and in accordance with the finding that estradiol benzoate (EB) treatment downregulated ERβ expression (Cao et al., 2012). ERα and ERβ bind the same consensus site and heterodimerize, suggesting that they can both compete and cooperate at the level of gene regulation (Bodo et al., 2006; Nilsson et al., 2001; Pettersson et al., 1997). This hypothesis is borne out at the behavioral level, as males mutant for both receptors show a more severe sexual behavior phenotype than the ERα mutants alone (Ogawa et al., 2000). ERβ upregulation in the BNST may therefore compensate for some aspects of organization mediated by ERα, which could account for the mild aggression phenotype in our *Vgat*-Cre mutant males. However, analysis of ERβ expression in mutant pups would be necessary to confirm if this upregulation occurs perinatally or after puberty.

ERβ has sexually dimorphic expression in the VMHvl with higher expression in females that is reduced to male levels by neonatal EB treatment (Ikeda et al., 2003). However, we do not detect any changes in Esr2 expression in the VMHvl of *Vglut2* mutant males (data not shown), or in the MePD of *Vgat* mutants (Figure 5J), suggesting that the Esr2 gene is regulated differently in the BNST compared to the MeA and VMHvl, both of which receive projections from the BNST. AR expression in VMHvl is also unaltered in *Vglut2* mutants (data not shown). We propose that loss of ERa in *Vgat+* neurons alters their function in male behaviors by demasculinizing gene expression. Future RNA-seq analysis of these populations would reveal the gene programs downstream of ERα that impart sex-specificity to neuronal function.

### Comparison of behavioral phenotypes in genetic and viral deletions of ERα

The behavioral alterations in our *Vgat* mutants are subtle compared to those seen in ERαKO males, particularly with regard to aggression. Although less than half of our mutants attack, this number does not reach significance when compared to all three control genotypes (Fig 4E, Table S1). Therefore the behavioral deficits from the deletion of *Esr1* in either excitatory or inhibitory neurons do not sum to the phenotype of the constitutive ERαKO males. We attribute these results to three primary factors. First, there are likely ERα+ cells that are not targeted by either *Vgat*-Cre or *Vglut2*-Cre, particularly in the MPOA (Fig 1F). Other neurotransmitters, such as dopamine, can be expressed in *Vgat/Vglut2*-negative cells, and may contribute important modulatory functions (Hnasko and Edwards, 2012). ERα is also expressed in non-neuronal cells such as astrocytes and endothelial cells (Arevalo et al., 2015; Azcoitia et al., 2010; Kuo et al., 2010) and these cell types could play a role in organizing and modulating behavioral circuitry. Second, our animals are on a mixed genetic background that is likely to obscure behavioral effects seen in a pure strain such as C57BL/6. Although both of our Cre lines were originally maintained on a mixed C57BL/6;FVB;129S6 genetic background (Vong et al., 2011), the *Vglut2*-Cre mice we received from Jackson possessed a white coat-color, while our founder *Vgat*-Cre mice were agouti (data not shown). Male sexual behaviors are highly dependent on genetic background (Dominguez-Salazar et al., 2004) and the initial characterization of gonadally intact ERαKO mutant males described much more robust mating behavior than was seen in later experiments when ERαKO mice were backcrossed to C57BL6/J (Ogawa et al., 1997; Wersinger and Rissman, 2000). We note that our three control genotypes display varying levels of male behaviors and serum testosterone levels, both within and between the *Vgat* and *Vglut2* cohorts. These results highlight the importance of testing all three control genotypes in this type of genetic cross. We therefore speculate that the behavior phenotype we see in our *Vgat*-Cre;*Esr1*^lox/lox^ males would be more severe if our experiments were performed in a pure C57BL6/J background. Finally, deletion of ERα in only a subset of the brain may allow for developmental compensation from other areas not targeted in each cross. We did not quantify ERα deletion in other ERα+ brain regions, such as the anteroventral periventricular nucleus (AVPV), arcuate nucleus, or paraventricular nucleus (PVN). This could be addressed by crossing the *Vglut2*-Cre and *Vgat*Cre-lines together to simultaneously delete *Esr1* in both excitatory and inhibitory neurons in the same animal.

Our behavioral results differ from those seen in a recent study in which acute knockdown of ERα via adenoviral delivery of shRNA into VMHvl decreased intensity of mating and aggression, although all knockdown males did intromit females and attack intruder males(Sano et al., 2013). These males also had increased body weight compared to shRNA controls, whereas our *Vglut2*-Cre;*Esr1*^lox/lox^ mutant males have similar weights to their control littermates (Fig 2A). It is possible that viral ERα knockdown males could recover their behaviors if tested at a later timepoint. Many of the available Cre drivers that target inhibitory neurons are tamoxifen inducible and therefore provide temporal control over deletion of a floxed allele (Taniguchi et al., 2011), although the timing of treatment with tamoxifen, an estrogen analog, needs careful consideration to avoid permanent disruption of sexual differentiation of the brain. It would be intriguing to delete ERα in the same population of neurons at different developmental timepoints to dissect out the distinct roles of this receptor in organizing or activating behavioral circuitry (McCarthy et al., 2009; Phoenix et al., 1959).

### The role of inhibitory neurons in male behavioral circuitry

Our results suggest that ERα masculinizes the brain by organizing inhibitory inputs onto glutamatergic neurons that drive behavioral output. This model is consistent with diverse previous studies on sexually dimorphic circuitry and behavior. First, there is extensive literature on the role of GABA in mediating sex differences in neuronal function (reviewed inMcCarthy et al., 2002). In particular, GABA signaling regulates the embryonic development of the VMH (Tobet et al., 2009). GABA actions on GABAB receptors are required for proper migration of ERα+ cells during VMH development (McClellan et al., 2008). Indeed, ERα itself is expressed in the developing VMH as early as e13.5, and is clearly defined in the VMHvl by e15.5 at which time GABA+ fibers already encircle the VMHvl (Tobet et al., 1999). It is likely that this innervation originates in ERα+ neurons from the BNST and MePD, as tracing studies in these regions show a similar pattern of afferents to the VMHvl in adult mice (Canteras et al., 1992; Choi et al., 2005; Dong and Swanson, 2004; Gu et al., 2003). Additionally, the VMHvl of males receives more input from aromatase-expressing neurons^33^. We propose that perinatal estrogen acts through ERα to masculinize this pre-existing circuit thereby leading to sex differences in VMHvl function in a cell non-autonomous fashion.

The primacy of inhibitory neurons in regulating sexually dimorphic behaviors has also been demonstrated in behavioral studies. Mice with genetic lesions to the VNO cannot detect non-volatile pheromonal cues from conspecifics and males display mating rather than aggression towards other males (Stowers et al., 2002). Therefore male olfactory cues initiate aggressive behavior by inhibiting the default mating behavioral repertoire, presumably through activation of GABAergic neurons in the MePD and BNST. Accordingly, moderate doses of GABA agonists elicit aggression behavior in rodents (Nelson and Trainor, 2007) and ablation of aromatase-expressing GABAergic neurons in the MePD attenuates aggression (Unger et al., 2015). Finally, optogenetic stimulation of *Vgat*+ MePD neurons drives time-locked mating and aggression behaviors in males (Hong et al., 2014) in a manner previously seen by stimulation of ERα/+*Vglut2*+ neurons in the VMHvl (Lee et al., 2014). As the MePD and BNST project strongly to each other and the VMHvl (Canteras et al., 1992; Choi et al., 2005; Dong and Swanson, 2004; Gu et al., 2003), it is likely that they cooperate to gate the disinhibition of the VMHvl, leading to the display of mating or aggression (Choi et al., 2005; Hong et al., 2014). We conclude that perinatal estrogen in the male brain sculpts this hierarchy of disinhibition through ERα-dependent gene expression programs in the MePD and BNST.

## Acknowledgements

We thank Sohaib Khan and Holly Ingraham for providing us with the Esr1^lox/lox^ mouse strain. We thank Stephen Shea for advice on statistics, and Stephanie Correa and Danielle Stolzenberg for comments on the manuscript. This work was performed with assistance from CSHL Shared Resources, including the Histology and Microscopy Core Facilities, which are supported by the Cancer Center Support Grant 5P30CA045508. This work was supported by a grant from Ted and Veda Stanley (Stanley Family Foundation).

---

**Figure S1:**
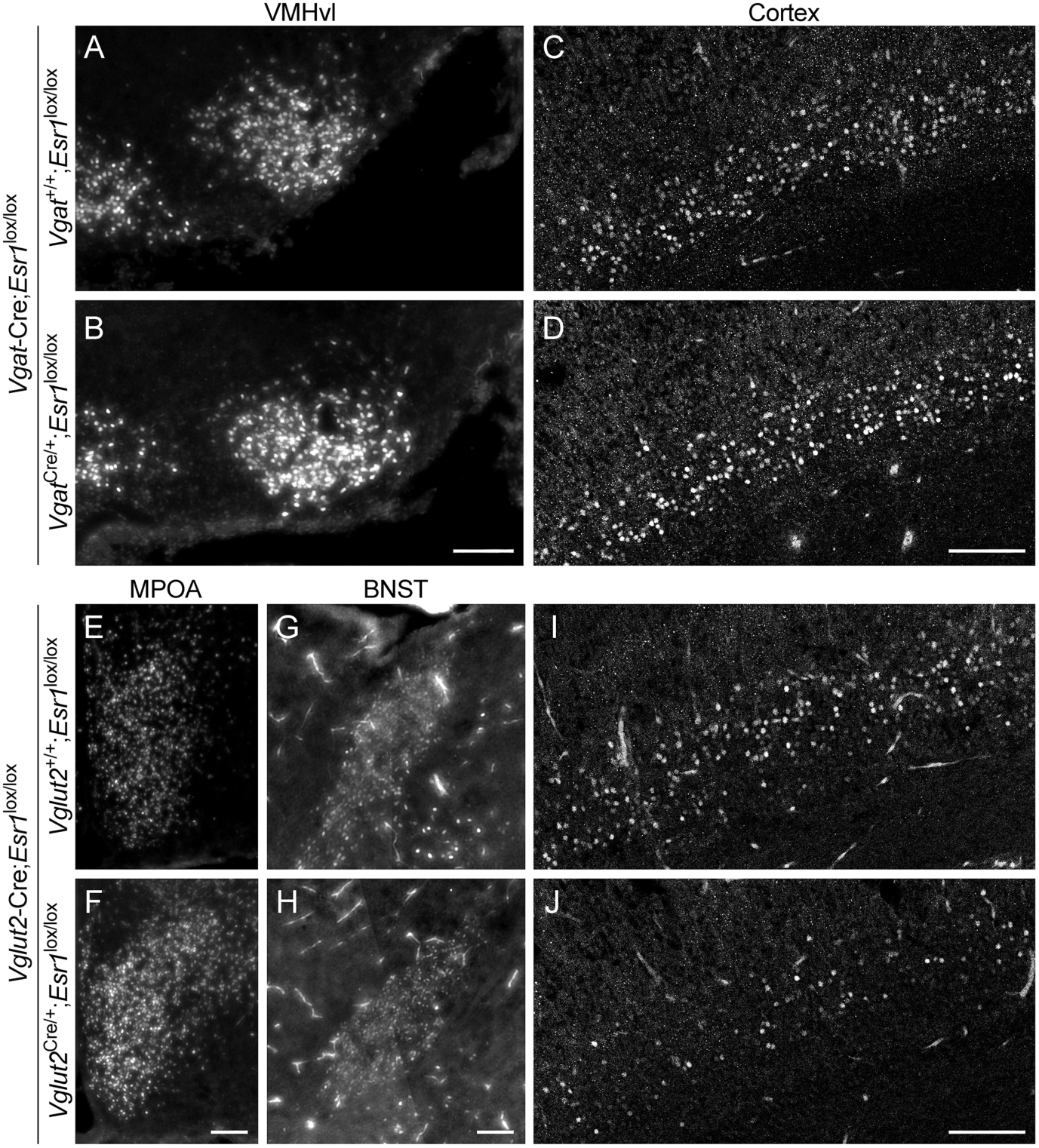
*Esr1* deletion is specific to the Cre-driver used. Representative images are shown of the VMHvl (**A,B**) and cortex (**C,D**) of lox/lox only control and *Vgat*-Cre;*Esr1*^lox/lox^ mutant littermates taken on p0. Also shown are the MPOA (**E,F**), BNST (**G,H**), and cortex (**I,J**) of lox/lox only and *Vglut2*-Cre;*Esr1*^lox/lox^ mutant p0 littermates. There is no apparent *Esr1* deletion in the VMHvl of *Vgat*-Cre, nor in the MPOA or BNST of *Vglut2*-Cre, mutant animals. ERα expression is reduced in the cortex of *Vglut2*-Cre but not *Vgat*-Cre mutant males. Scalebars = 100μ.

**Figure S2:**
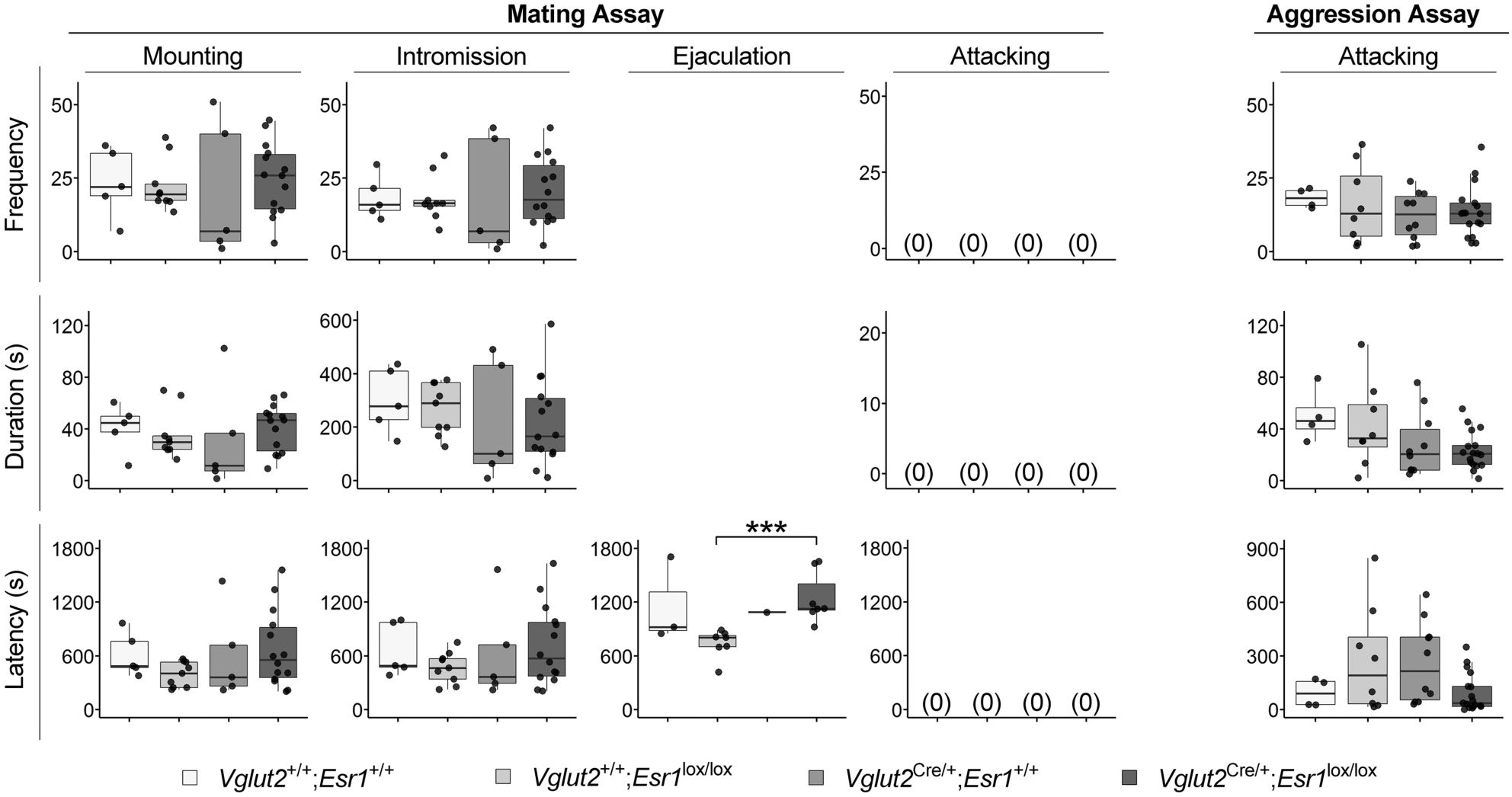
No qualitative differences in sex-specific behaviors exhibited by *Vglut2*-Cre;*Esr1*^lox/lox^ mutant animals. The total frequency, total duration, and latency to first instance of mounting, intromission, ejaculation, and attacking behavior in mating assays as well as attacking behavior in aggression assays is depicted. Boxplots denote median and 1^st^ and 3^rd^ quartiles. Whiskers denote 1.5^*^ interquartile range. ***p<0.005, Kruskal Wallis omnibus test followed by Dunn’s post-hoc test for multiple comparisons with one control.

**Figure S3:**
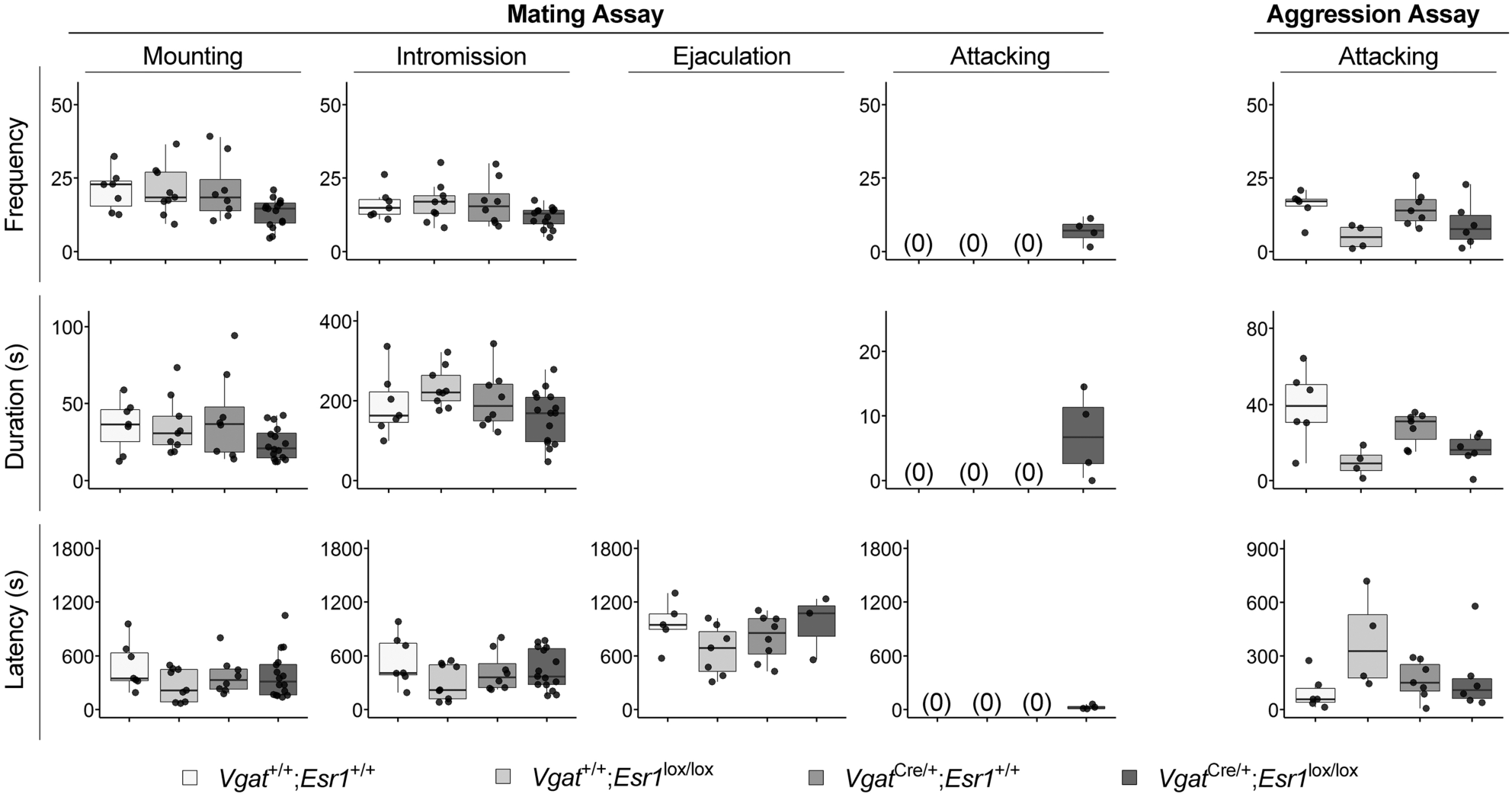
*Vgat*-Cre;*Esr1*^lox/lox^ mutant animals exhibit no qualitative deficits in mating and aggressive behavior. The total frequency, total duration, and latency to first instance of mounting, intromission, ejaculation, and attacking behavior in mating assays as well as attacking behavior in aggression assays is depicted. Boxplots denote median and 1^st^ and 3^rd^ quartiles. Whiskers denote 1.5^*^ interquartile range.

**Table S1:** Detailed data and statistical analyses. All data and statistical tests as shown in each figure are provided in detail. IQR denotes inter-quartile range between 1^st^ and 3^rd^ quartiles. Italicized p-values are assigned significance.

Statistics for Figure 1
| Region | Genotype | Median | IQR | n | Mann Whitney<br>U Statistic | p-value |
| --- | --- | --- | --- | --- | --- | --- |
| VMHvl | *Vglut2*<sup>+/+</sup>;*Esr1*<sup>lox/lox</sup> | 4808.0 | 1254.5 | 4 | 16 | *0.021* |
|  | *Vglut2*<sup>Cre/+</sup>;*Esr1*<sup>lox/lox</sup> | 69.0 | 30.5 | 4 |  |  |
| MePV | *Vglut2*<sup>+/+</sup>;*Esr1*<sup>lox/lox</sup> | 2372.0 | 621.0 | 4 | 16 | *0.021* |
|  | *Vglut2*<sup>Cre/+</sup>;*Esr1*<sup>lox/lox</sup> | 90.0 | 44.0 | 4 |  |  |
| MPOA | *Vgat*<sup>+/+</sup>;*Esr1*<sup>lox/lox</sup> | 9879.0 | 3963.0 | 4 | 16 | *0.021* |
|  | *Vgat*<sup>Cre/+</sup>;*Esr1*<sup>lox/lox</sup> | 1639.0 | 780.5 | 4 |  |  |
| BNST | *Vgat*<sup>+/+</sup>;*Esr1*<sup>lox/lox</sup> | 2899.0 | 601.5 | 4 | 16 | *0.021* |
|  | *Vgat*<sup>Cre/+</sup>;*Esr1*<sup>lox/lox</sup> | 80.0 | 109.0 | 4 |  |  |
| MePD | *Vgat*<sup>+/+</sup>;*Esr1*<sup>lox/lox</sup> | 10061.0 | 1618.5 | 4 | 16 | *0.021* |
|  | *Vgat*<sup>Cre/+</sup>;*Esr1*<sup>lox/lox</sup> | 536.0 | 69.0 | 4 |  |  |

Statistics for Figure 2
| Measurement | Genotype | Median | IQR | n | Kruskal Wallis H statistic | p-value | Dunn's post-hoc against mutant |
| --- | --- | --- | --- | --- | --- | --- | --- |
| Body weight (g) | *Vglut2*<sup>+/+</sup>;*Esr1*<sup>+/+</sup> | 31.3 | 0.8 | 5 | 2.78 | 0.426 |  |
|  | *Vglut2*<sup>+/+</sup>;*Esr1*<sup>lox/lox</sup> | 28.8 | 2.6 | 10 |  |  |  |
|  | *Vglut2*<sup>Cre/+</sup>;*Esr1*<sup>+/+</sup> | 29.6 | 2.4 | 10 |  |  |  |
|  | *Vglut2*<sup>Cre/+</sup>;*Esr1*<sup>lox/lox</sup> | 30.0 | 4.5 | 20 |  |  |  |
|  | *Vgat*<sup>+/+</sup>;*Esr1*<sup>+/+</sup> | 29.2 | 2.5 | 7 | 6.77 | 0.080 |  |
|  | *Vgat*<sup>+/+</sup>;*Esr1*<sup>lox/lox</sup> | 30.5 | 2.2 | 9 |  |  |  |
|  | *Vgat*<sup>Cre/+</sup>;*Esr1*<sup>+/+</sup> | 31.2 | 4.4 | 8 |  |  |  |
|  | *Vgat*<sup>Cre/+</sup>;*Esr1*<sup>lox/lox</sup> | 29.2 | 2.9 | 16 |  |  |  |
| Testes weight (g) | *Vglut2*<sup>+/+</sup>;*Esr1*<sup>+/+</sup> | 0.213 | 0.008 | 5 | 3.27 | 0.352 |  |
|  | *Vglut2*<sup>+/+</sup>;*Esr1*<sup>lox/lox</sup> | 0.202 | 0.022 | 10 |  |  |  |
|  | *Vglut2*<sup>Cre/+</sup>;*Esr1*<sup>+/+</sup> | 0.194 | 0.027 | 10 |  |  |  |
|  | *Vglut2*<sup>Cre/+</sup>;*Esr1*<sup>lox/lox</sup> | 0.213 | 0.031 | 20 |  |  |  |
|  | *Vgat*<sup>+/+</sup>;*Esr1*<sup>+/+</sup> | 0.176 | 0.031 | 7 | 5.76 | 0.124 |  |
|  | *Vgat*<sup>+/+</sup>;*Esr1*<sup>lox/lox</sup> | 0.194 | 0.017 | 9 |  |  |  |
|  | *Vgat*<sup>Cre/+</sup>;*Esr1*<sup>+/+</sup> | 0.178 | 0.021 | 8 |  |  |  |
|  | *Vgat*<sup>Cre/+</sup>;*Esr1*<sup>lox/lox</sup> | 0.209 | 0.037 | 16 |  |  |  |
| Serum testosterone (nM) | *Vglut2*<sup>+/+</sup>;*Esr1*<sup>+/+</sup> | 1.86 | 3.22 | 5 | 15.68 | 0.0013 | 0.045 |
|  | *Vglut2*<sup>+/+</sup>;*Esr1*<sup>lox/lox</sup> | 3.35 | 2.65 | 10 |  |  | 0.077 |
|  | *Vglut2*<sup>Cre/+</sup>;*Esr1*<sup>+/+</sup> | 2.49 | 1.10 | 10 |  |  | 0.001 |
|  | *Vglut2*<sup>Cre/+</sup>;*Esr1*<sup>lox/lox</sup> | 19.78 | 22.22 | 20 |  |  |  |
|  | *Vgat*<sup>+/+</sup>;*Esr1*<sup>+/+</sup> | 2.32 | 3.24 | 7 | 3.27 | 0.352 |  |
|  | *Vgat*<sup>+/+</sup>;*Esr1*<sup>lox/lox</sup> | 8.92 | 16.32 | 9 |  |  |  |
|  | *Vgat*<sup>Cre/+</sup>;*Esr1*<sup>+/+</sup> | 16.83 | 37.68 | 8 |  |  |  |
|  | *Vgat*<sup>Cre/+</sup>;*Esr1*<sup>lox/lox</sup> | 2.83 | 8.89 | 16 |  |  |  |
| Seminal vesicles weight (g) | *Vglut2*<sup>+/+</sup>;*Esr1*<sup>+/+</sup> | 0.304 | 0.083 | 5 | 8.60 | 0.035 | 0.991 |
|  | *Vglut2*<sup>+/+</sup>;*Esr1*<sup>lox/lox</sup> | 0.292 | 0.125 | 10 |  |  | 0.068 |
|  | *Vglut2*<sup>Cre/+</sup>;*Esr1*<sup>+/+</sup> | 0.267 | 0.028 | 10 |  |  | 0.035 |
|  | *Vglut2*<sup>Cre/+</sup>;*Esr1*<sup>lox/lox</sup> | 0.368 | 0.155 | 20 |  |  |  |
|  | *Vgat*<sup>+/+</sup>;*Esr1*<sup>+/+</sup> | 0.240 | 0.039 | 7 | 9.48 | 0.024 | 0.030 |
|  | *Vgat*<sup>+/+</sup>;*Esr1*<sup>lox/lox</sup> | 0.291 | 0.040 | 9 |  |  | 1.000 |
|  | *Vgat*<sup>Cre/+</sup>;*Esr1*<sup>+/+</sup> | 0.257 | 0.063 | 8 |  |  | 0.180 |
|  | *Vgat*<sup>Cre/+</sup>;*Esr1*<sup>lox/lox</sup> | 0.292 | 0.039 | 16 |  |  |  |

Statistics for Figure 3
| Behavior | Genotype | % Animals performing | n | Fisher's 2x4 p-value | post-hoc Fisher's 2x2 against mutant |
| --- | --- | --- | --- | --- | --- |
| Mating--Mounting | *Vglut2*<sup>+/+</sup>;*Esr1*<sup>+/+</sup> | 100% | 5 | 0.135 |  |
|  | *Vglut2*<sup>+/+</sup>;*Esr1*<sup>lox/lox</sup> | 90% | 10 |  |  |
|  | *Vglut2*<sup>Cre/+</sup>;*Esr1*<sup>+/+</sup> | 50% | 10 |  |  |
|  | *Vglut2*<sup>Cre/+</sup>;*Esr1*<sup>lox/lox</sup> | 70% | 20 |  |  |
| Mating--Intromission | *Vglut2*<sup>+/+</sup>;*Esr1*<sup>+/+</sup> | 100% | 5 | 0.135 |  |
|  | *Vglut2*<sup>+/+</sup>;*Esr1*<sup>lox/lox</sup> | 90% | 10 |  |  |
|  | *Vglut2*<sup>Cre/+</sup>;*Esr1*<sup>+/+</sup> | 50% | 10 |  |  |
|  | *Vglut2*<sup>Cre/+</sup>;*Esr1*<sup>lox/lox</sup> | 70% | 20 |  |  |
| Mating--Ejaculation | *Vglut2*<sup>+/+</sup>;*Esr1*<sup>+/+</sup> | 60% | 5 | 0.031 | 0.122 |
|  | *Vglut2*<sup>+/+</sup>;*Esr1*<sup>lox/lox</sup> | 70% | 10 |  | 0.210 |
|  | *Vglut2*<sup>Cre/+</sup>;*Esr1*<sup>+/+</sup> | 10% | 10 |  | 0.358 |
|  | *Vglut2*<sup>Cre/+</sup>;*Esr1*<sup>lox/lox</sup> | 35% | 20 |  |  |
| Mating--Attacking | *Vglut2*<sup>+/+</sup>;*Esr1*<sup>+/+</sup> | 0% | 5 | 1.000 |  |
|  | *Vglut2*<sup>+/+</sup>;*Esr1*<sup>lox/lox</sup> | 0% | 10 |  |  |
|  | *Vglut2*<sup>Cre/+</sup>;*Esr1*<sup>+/+</sup> | 0% | 10 |  |  |
|  | *Vglut2*<sup>Cre/+</sup>;*Esr1*<sup>lox/lox</sup> | 0% | 20 |  |  |
| Aggression--Attacking | *Vglut2*<sup>+/+</sup>;*Esr1*<sup>+/+</sup> | 80% | 5 | 0.592 |  |
|  | *Vglut2*<sup>+/+</sup>;*Esr1*<sup>lox/lox</sup> | 80% | 10 |  |  |
|  | *Vglut2*<sup>Cre/+</sup>;*Esr1*<sup>+/+</sup> | 100% | 10 |  |  |
|  | *Vglut2*<sup>Cre/+</sup>;*Esr1*<sup>lox/lox</sup> | 85% | 20 |  |  |

| Measurement | Genotype | Median | IQR | Kruskal Wallis H statistic | p-value |
| --- | --- | --- | --- | --- | --- |
| # Spots | *Vglut2*<sup>+/+</sup>;*Esr1*<sup>+/+</sup> | 125.0 | 200.0 | 2.0163 | 0.569 |
|  | *Vglut2*<sup>+/+</sup>;*Esr1*<sup>lox/lox</sup> | 261.0 | 285.3 |  |  |
|  | *Vglut2*<sup>Cre/+</sup>;*Esr1*<sup>+/+</sup> | 268.5 | 105.5 |  |  |
|  | *Vglut2*<sup>Cre/+</sup>;*Esr1*<sup>lox/lox</sup> | 222.5 | 301.3 |  |  |
| Total Spot Area (cm<sup>2</sup>) | *Vglut2*<sup>+/+</sup>;*Esr1*<sup>+/+</sup> | 43.3 | 29.4 | 7.4104 | 0.060 |
|  | *Vglut2*<sup>+/+</sup>;*Esr1*<sup>lox/lox</sup> | 64.8 | 20.0 |  |  |
|  | *Vglut2*<sup>Cre/+</sup>;*Esr1*<sup>+/+</sup> | 68.5 | 27.8 |  |  |
|  | *Vglut2*<sup>Cre/+</sup>;*Esr1*<sup>lox/lox</sup> | 47.0 | 36.3 |  |  |

Statistics for Figure 4
| Behavior | Genotype | % Animals performing | n | Fisher's 2x4 p-value | post-hoc Fisher's 2x2 against mutant |
| --- | --- | --- | --- | --- | --- |
| Mating--<br>Mounting | *Vgat*<sup>+/+</sup>;*Esr1*<sup>+/+</sup> | 100% | 7 | 1.000 |  |
|  | *Vgat*<sup>+/+</sup>;*Esr1*<sup>lox/lox</sup> | 100% | 9 |  |  |
|  | *Vgat*<sup>Cre/+</sup>;*Esr1*<sup>+/+</sup> | 100% | 8 |  |  |
|  | *Vgat*<sup>Cre/+</sup>;*Esr1*<sup>lox/lox</sup> | 100% | 16 |  |  |
| Mating--<br>Intromission | *Vgat*<sup>+/+</sup>;*Esr1*<sup>+/+</sup> | 100% | 7 | 1.000 |  |
|  | *Vgat*<sup>+/+</sup>;*Esr1*<sup>lox/lox</sup> | 100% | 9 |  |  |
|  | *Vgat*<sup>Cre/+</sup>;*Esr1*<sup>+/+</sup> | 100% | 8 |  |  |
|  | *Vgat*<sup>Cre/+</sup>;*Esr1*<sup>lox/lox</sup> | 94% | 16 |  |  |
| Mating--<br>Ejaculation | *Vgat*<sup>+/+</sup>;*Esr1*<sup>+/+</sup> | 72% | 7 | 0.0002 | 0.026 |
|  | *Vgat*<sup>+/+</sup>;*Esr1*<sup>lox/lox</sup> | 78% | 9 |  | 0.009 |
|  | *Vgat*<sup>Cre/+</sup>;*Esr1*<sup>+/+</sup> | 100% | 8 |  | 0.0002 |
|  | *Vgat*<sup>Cre/+</sup>;*Esr1*<sup>lox/lox</sup> | 19% | 16 |  |  |
| Mating--<br>Attacking | *Vgat*<sup>+/+</sup>;*Esr1*<sup>+/+</sup> | 0% | 7 | 0.145 |  |
|  | *Vgat*<sup>+/+</sup>;*Esr1*<sup>lox/lox</sup> | 0% | 9 |  |  |
|  | *Vgat*<sup>Cre/+</sup>;*Esr1*<sup>+/+</sup> | 0% | 8 |  |  |
|  | *Vgat*<sup>Cre/+</sup>;*Esr1*<sup>lox/lox</sup> | 25% | 16 |  |  |
| Aggression--<br>Attacking | *Vgat*<sup>+/+</sup>;*Esr1*<sup>+/+</sup> | 86% | 7 | 0.040 | 0.069 |
|  | *Vgat*<sup>+/+</sup>;*Esr1*<sup>lox/lox</sup> | 44% | 9 |  | 1.000 |
|  | *Vgat*<sup>Cre/+</sup>;*Esr1*<sup>+/+</sup> | 88% | 8 |  | 0.033 |
|  | *Vgat*<sup>Cre/+</sup>;*Esr1*<sup>lox/lox</sup> | 38% | 16 |  |  |

| Measurement | Genotype | Median | IQR | Kruskal Wallis H statistic | p-value | Dunn's post-hoc against mutant |
| --- | --- | --- | --- | --- | --- | --- |
| # Spots | *Vgat*<sup>+/+</sup>;*Esr1*<sup>+/+</sup> | 108.0 | 74.5 | 21.966 | 0.00007 | 0.037 |
|  | *Vgat*<sup>+/+</sup>;*Esr1*<sup>lox/lox</sup> | 168.0 | 186.0 |  |  | 0.0006 |
|  | *Vgat*<sup>Cre/+</sup>;*Esr1*<sup>+/+</sup> | 195.5 | 81.5 |  |  | 0.0003 |
|  | *Vgat*<sup>Cre/+</sup>;*Esr1*<sup>lox/lox</sup> | 49.0 | 39.3 |  |  |  |
| Total Spot Area (cm<sup>2</sup>) | *Vgat*<sup>+/+</sup>;*Esr1*<sup>+/+</sup> | 74.3 | 40.6 | 20.446 | 0.00014 | 0.008 |
|  | *Vgat*<sup>+/+</sup>;*Esr1*<sup>lox/lox</sup> | 56.7 | 8.3 |  |  | 0.012 |
|  | *Vgat*<sup>Cre/+</sup>;*Esr1*<sup>+/+</sup> | 90.4 | 24.1 |  |  | 0.0002 |
|  | *Vgat*<sup>Cre/+</sup>;*Esr1*<sup>lox/lox</sup> | 35.1 | 18.3 |  |  |  |

Statistics for Figure 5
| Gene | Region | Genotype | Median | IQR | n | Mann Whitney<br>U Statistic | p-value |
| --- | --- | --- | --- | --- | --- | --- | --- |
| AR | BNST | *Vgat*<sup>+/+</sup>;*Esr1*<sup>lox/lox</sup> | 189.2 | 35.1 | 4 | 15 | 0.043 |
|  |  | *Vgat*<sup>Cre/+</sup>;*Esr1*<sup>lox/lox</sup> | 103.8 | 24.4 | 4 |  |  |
|  | MePD | *Vgat*<sup>+/+</sup>;*Esr1*<sup>lox/lox</sup> | 265.9 | 40.5 | 4 | 15 | 0.043 |
|  |  | *Vgat*<sup>Cre/+</sup>;*Esr1*<sup>lox/lox</sup> | 154.0 | 41.8 | 4 |  |  |
| Esr2 | BNST | *Vgat*<sup>+/+</sup>;*Esr1*<sup>lox/lox</sup> | 58.4 | 7.6 | 4 | 15 | 0.043 |
|  |  | *Vgat*<sup>Cre/+</sup>;*Esr1*<sup>lox/lox</sup> | 75.1 | 2.0 | 4 |  |  |
|  | MePD | *Vgat*<sup>+/+</sup>;*Esr1*<sup>lox/lox</sup> | 86.5 | 32.8 | 4 | 8 | >0.999 |
|  |  | *Vgat*<sup>Cre/+</sup>;*Esr1*<sup>lox/lox</sup> | 114.4 | 68.0 | 4 |  |  |

Statistics for Supplementary Figure S2
| Behavior | Parameter | Genotype | Median | IQR | Kruskal Wallis H statistic | p-value | Dunn's post-hoc against mutant |
| --- | --- | --- | --- | --- | --- | --- | --- |
| Mating--Mounting | Frequency | *Vglut2*<sup>+/+</sup>;*Esr1*<sup>+/+</sup> | 22.00 | 14.50 | 0.61 | 0.895 |  |
|  |  | *Vglut2*<sup>+/+</sup>;*Esr1*<sup>lox/lox</sup> | 19.50 | 5.50 |  |  |  |
|  |  | *Vglut2*<sup>Cre/+</sup>;*Esr1*<sup>+/+</sup> | 7.00 | 36.50 |  |  |  |
|  |  | *Vglut2*<sup>Cre/+</sup>;*Esr1*<sup>lox/lox</sup> | 26.00 | 18.50 |  |  |  |
|  | Duration (s) | *Vglut2*<sup>+/+</sup>;*Esr1*<sup>+/+</sup> | 44.63 | 12.25 | 2.27 | 0.519 |  |
|  |  | *Vglut2*<sup>+/+</sup>;*Esr1*<sup>lox/lox</sup> | 29.88 | 10.46 |  |  |  |
|  |  | *Vglut2*<sup>Cre/+</sup>;*Esr1*<sup>+/+</sup> | 11.68 | 29.20 |  |  |  |
|  |  | *Vglut2*<sup>Cre/+</sup>;*Esr1*<sup>lox/lox</sup> | 46.67 | 29.16 |  |  |  |
|  | Latency (s) | *Vglut2*<sup>+/+</sup>;*Esr1*<sup>+/+</sup> | 486.04 | 290.95 | 3.04 | 0.386 |  |
|  |  | *Vglut2*<sup>+/+</sup>;*Esr1*<sup>lox/lox</sup> | 404.62 | 282.07 |  |  |  |
|  |  | *Vglut2*<sup>Cre/+</sup>;*Esr1*<sup>+/+</sup> | 362.86 | 458.32 |  |  |  |
|  |  | *Vglut2*<sup>Cre/+</sup>;*Esr1*<sup>lox/lox</sup> | 555.69 | 558.57 |  |  |  |
| Mating--Intromission | Frequency | *Vglut2*<sup>+/+</sup>;*Esr1*<sup>+/+</sup> | 16.00 | 7.50 | 0.48 | 0.923 |  |
|  |  | *Vglut2*<sup>+/+</sup>;*Esr1*<sup>lox/lox</sup> | 16.50 | 2.00 |  |  |  |
|  |  | *Vglut2*<sup>Cre/+</sup>;*Esr1*<sup>+/+</sup> | 7.00 | 35.50 |  |  |  |
|  |  | *Vglut2*<sup>Cre/+</sup>;*Esr1*<sup>lox/lox</sup> | 17.75 | 18.00 |  |  |  |
|  | Duration (s) | *Vglut2*<sup>+/+</sup>;*Esr1*<sup>+/+</sup> | 279.09 | 182.00 | 2.51 | 0.474 |  |
|  |  | *Vglut2*<sup>+/+</sup>;*Esr1*<sup>lox/lox</sup> | 290.24 | 166.13 |  |  |  |
|  |  | *Vglut2*<sup>Cre/+</sup>;*Esr1*<sup>+/+</sup> | 100.56 | 366.77 |  |  |  |
|  |  | *Vglut2*<sup>Cre/+</sup>;*Esr1*<sup>lox/lox</sup> | 166.22 | 197.31 |  |  |  |
|  | Latency (s) | *Vglut2*<sup>+/+</sup>;*Esr1*<sup>+/+</sup> | 489.23 | 497.29 | 1.64 | 0.650 |  |
|  |  | *Vglut2*<sup>+/+</sup>;*Esr1*<sup>lox/lox</sup> | 466.71 | 225.47 |  |  |  |
|  |  | *Vglut2*<sup>Cre/+</sup>;*Esr1*<sup>+/+</sup> | 366.43 | 428.43 |  |  |  |
|  |  | *Vglut2*<sup>Cre/+</sup>;*Esr1*<sup>lox/lox</sup> | 571.66 | 595.74 |  |  |  |
| Mating—Ejaculation | Latency (s) | *Vglut2*<sup>+/+</sup>;*Esr1*<sup>+/+</sup> | 919.69 | 428.81 | 12.15 | 0.007 | 1.000 |
|  |  | *Vglut2*<sup>+/+</sup>;*Esr1*<sup>lox/lox</sup> | 805.21 | 125.64 |  |  | 0.002 |
|  |  | *Vglut2*<sup>Cre/+</sup>;*Esr1*<sup>+/+</sup> | 1087.15 | 0.00 |  |  | 1.000 |
|  |  | *Vglut2*<sup>Cre/+</sup>;*Esr1*<sup>lox/lox</sup> | 1127.32 | 296.59 |  |  |  |
| Aggression--Attacking | Frequency | *Vglut2*<sup>+/+</sup>;*Esr1*<sup>+/+</sup> | 18.25 | 5.00 | 1.99 | 0.574 |  |
|  |  | *Vglut2*<sup>+/+</sup>;*Esr1*<sup>lox/lox</sup> | 13.00 | 20.50 |  |  |  |
|  |  | *Vglut2*<sup>Cre/+</sup>;*Esr1*<sup>+/+</sup> | 12.75 | 13.00 |  |  |  |
|  |  | *Vglut2*<sup>Cre/+</sup>;*Esr1*<sup>lox/lox</sup> | 13.00 | 7.00 |  |  |  |
|  | Duration<br>(s) | *Vglut2*<sup>+/+</sup>;*Esr1*<sup>+/+</sup> | 46.08 | 16.27 | 6.41 | 0.093 |  |
|  |  | *Vglut2*<sup>+/+</sup>;*Esr1*<sup>lox/lox</sup> | 32.73 | 32.62 |  |  |  |
|  |  | *Vglut2*<sup>Cre/+</sup>;*Esr1*<sup>+/+</sup> | 20.66 | 31.58 |  |  |  |
|  |  | *Vglut2*<sup>Cre/+</sup>;*Esr1*<sup>lox/lox</sup> | 20.97 | 14.65 |  |  |  |
|  | Latency (s) | *Vglut2*<sup>+/+</sup>;*Esr1*<sup>+/+</sup> | 90.21 | 129.31 | 6.71 | 0.082 |  |
|  |  | *Vglut2*<sup>+/+</sup>;*Esr1*<sup>lox/lox</sup> | 192.83 | 373.30 |  |  |  |
|  |  | *Vglut2*<sup>Cre/+</sup>;*Esr1*<sup>+/+</sup> | 216.87 | 350.26 |  |  |  |
|  |  | *Vglut2*<sup>Cre/+</sup>;*Esr1*<sup>lox/lox</sup> | 37.18 | 111.77 |  |  |  |

Statistics for Supplementary Figure S3
| Behavior | Parameter | Genotype | Median | IQR | Kruskal Wallis H statistic | p-value | Dunn's post-hoc against mutant |
| --- | --- | --- | --- | --- | --- | --- | --- |
| Mating--Mounting | Frequency | *Vgat*<sup>+/+</sup>;*Esr1*<sup>+/+</sup> | 23.00 | 8.50 | 8.89 | 0.031 | 0.057 |
|  |  | *Vgat*<sup>+/+</sup>;*Esr1*<sup>lox/lox</sup> | 18.50 | 10.00 |  |  | 0.065 |
|  |  | *Vgat*<sup>Cre/+</sup>;*Esr1*<sup>+/+</sup> | 18.50 | 10.63 |  |  | 0.144 |
|  |  | *Vgat*<sup>Cre/+</sup>;*Esr1*<sup>lox/lox</sup> | 14.75 | 6.75 |  |  |  |
|  | Duration (s) | *Vgat*<sup>+/+</sup>;*Esr1*<sup>+/+</sup> | 36.67 | 20.75 | 4.65 | 0.199 |  |
|  |  | *Vgat*<sup>+/+</sup>;*Esr1*<sup>lox/lox</sup> | 30.75 | 18.67 |  |  |  |
|  |  | *Vgat*<sup>Cre/+</sup>;*Esr1*<sup>+/+</sup> | 36.87 | 29.50 |  |  |  |
|  |  | *Vgat*<sup>Cre/+</sup>;*Esr1*<sup>lox/lox</sup> | 20.97 | 16.15 |  |  |  |
|  | Latency (s) | *Vgat*<sup>+/+</sup>;*Esr1*<sup>+/+</sup> | 349.79 | 307.17 | 2.73 | 0.435 |  |
|  |  | *Vgat*<sup>+/+</sup>;*Esr1*<sup>lox/lox</sup> | 217.20 | 362.88 |  |  |  |
|  |  | *Vgat*<sup>Cre/+</sup>;*Esr1*<sup>+/+</sup> | 332.00 | 225.38 |  |  |  |
|  |  | *Vgat*<sup>Cre/+</sup>;*Esr1*<sup>lox/lox</sup> | 313.44 | 340.82 |  |  |  |
| Mating--Intromission | Frequency | *Vgat*<sup>+/+</sup>;*Esr1*<sup>+/+</sup> | 15.00 | 5.00 | 4.83 | 0.185 |  |
|  |  | *Vgat*<sup>+/+</sup>;*Esr1*<sup>lox/lox</sup> | 17.00 | 6.00 |  |  |  |
|  |  | *Vgat*<sup>Cre/+</sup>;*Esr1*<sup>+/+</sup> | 15.50 | 9.25 |  |  |  |
|  |  | *Vgat*<sup>Cre/+</sup>;*Esr1*<sup>lox/lox</sup> | 13.00 | 4.50 |  |  |  |
|  | Duration (s) | *Vgat*<sup>+/+</sup>;*Esr1*<sup>+/+</sup> | 163.17 | 77.21 | 6.53 | 0.089 |  |
|  |  | *Vgat*<sup>+/+</sup>;*Esr1*<sup>lox/lox</sup> | 221.05 | 63.90 |  |  |  |
|  |  | *Vgat*<sup>Cre/+</sup>;*Esr1*<sup>+/+</sup> | 187.46 | 90.99 |  |  |  |
|  |  | *Vgat*<sup>Cre/+</sup>;*Esr1*<sup>lox/lox</sup> | 169.33 | 111.28 |  |  |  |
|  | Latency (s) | *Vgat*<sup>+/+</sup>;*Esr1*<sup>+/+</sup> | 410.51 | 354.45 | 3.63 | 0.305 |  |
|  |  | *Vgat*<sup>+/+</sup>;*Esr1*<sup>lox/lox</sup> | 218.48 | 378.34 |  |  |  |
|  |  | *Vgat*<sup>Cre/+</sup>;*Esr1*<sup>+/+</sup> | 362.15 | 267.04 |  |  |  |
|  |  | *Vgat*<sup>Cre/+</sup>;*Esr1*<sup>lox/lox</sup> | 371.36 | 400.73 |  |  |  |
| Mating—Ejaculation | Latency (s) | *Vgat*<sup>+/+</sup>;*Esr1*<sup>+/+</sup> | 948.76 | 173.07 | 4.12 | 0.249 |  |
|  |  | *Vgat*<sup>+/+</sup>;*Esr1*<sup>lox/lox</sup> | 689.00 | 443.68 |  |  |  |
|  |  | *Vgat*<sup>Cre/+</sup>;*Esr1*<sup>+/+</sup> | 854.81 | 395.36 |  |  |  |
|  |  | *Vgat*<sup>Cre/+</sup>;*Esr1*<sup>lox/lox</sup> | 1075.64 | 339.40 |  |  |  |
| Mating—Attacking | Frequency | *Vgat*<sup>Cre/+</sup>;*Esr1*<sup>lox/lox</sup> | 7.25 | 4.63 |  |  |  |
|  | Duration (s) | *Vgat*<sup>Cre/+</sup>;*Esr1*<sup>lox/lox</sup> | 6.75 | 8.75 |  |  |  |
|  | Latency (s) | *Vgat*<sup>Cre/+</sup>;*Esr1*<sup>lox/lox</sup> | 18.29 | 21.36 |  |  |  |
| Aggression--Attacking | Frequency | $Vgat^{+/+};Esr1^{+/+}$ | 17.25 | 2.38 | 7.98 | 0.046 | 0.280 |
| | | $Vgat^{+/+};Esr1^{lox/lox}$ | 5.00 | 6.50 | | | 1.000 |
| | | $Vgat^{Cre/+};Esr1^{+/+}$ | 14.00 | 7.25 | | | 0.410 |
| | | $Vgat^{Cre/+};Esr1^{lox/lox}$ | 7.75 | 8.13 | | | |
| | Duration<br>(s) | $Vgat^{+/+};Esr1^{+/+}$ | 39.31 | 20.06 | 10.30 | 0.016 | 0.081 |
| | | $Vgat^{+/+};Esr1^{lox/lox}$ | 9.09 | 8.17 | | | 1.000 |
| | | $Vgat^{Cre/+};Esr1^{+/+}$ | 31.16 | 11.97 | | | 0.210 |
| | | $Vgat^{Cre/+};Esr1^{lox/lox}$ | 16.15 | 8.07 | | | |
| | Latency (s) | $Vgat^{+/+};Esr1^{+/+}$ | 58.18 | 76.72 | 5.69 | 0.128 | |
| | | $Vgat^{+/+};Esr1^{lox/lox}$ | 328.48 | 353.34 | | | |
| | | $Vgat^{Cre/+};Esr1^{+/+}$ | 150.97 | 149.01 | | | |
| | | $Vgat^{Cre/+};Esr1^{lox/lox}$ | 109.43 | 111.52 | | | |

